# CIDANE: Comprehensive isoform discovery and abundance estimation

**DOI:** 10.1101/017939

**Authors:** Stefan Canzar, Sandro Andreotti, David Weese, Knut Reinert, Gunnar W. Klau

**Author notes:** Equal contribution (Stefan Canzar, Sandro Andreotti). Shared last author (Knut Reinert, Gunnar W. Klau).

## Abstract

We present CIDANE, a novel framework for genome-based transcript reconstruction and quantification from RNA-seq reads. CIDANE assembles transcripts with significantly higher sensitivity and precision than existing tools, while competing in speed with the fastest methods. In addition to reconstructing transcripts *ab initio*, the algorithm also allows to make use of the growing annotation of known splice sites, transcription start and end sites, or full-length transcripts, which are available for most model organisms. CIDANE supports the integrated analysis of RNA-seq and additional gene-boundary data and recovers splice junctions that are invisible to other methods. CIDANE is available at http://ccb.jhu.edu/software/cidane/.

## 1 Background

High-throughput sequencing of cellular RNA (RNA-seq) aims at identifying and quantifying the set of all RNA molecules, the transcriptome, produced by a cell. Despite having largely identical genomes, the RNA content of cells differs among tissues, developmental stages, and between disease and normal condition. For eukaryotes, differences are determined by the set of genes being expressed, but also by the different mRNA isoforms each gene may produce; alternative splicing and alternative transcription and polyadenylation define and combine exons in distinct ways.

RNA-seq technology can generate hundreds of millions of short (50-250bp) strings of bases, called reads, from expressed transcripts at a fraction of the time and cost required by conventional Sanger sequencing. The wealth of RNA-seq data produced recently has revealed novel isoforms [1, 2, 3] and new classes of RNA [4], allowed a better characterization of cancer transcriptomes [5, 6], and led to the discovery of splicing aberrations in disease [7, 8].

However, the step from sequencing to profiling the cellular transcriptome involves solving a high-dimensional, complex puzzle, which poses major challenges to bioinformatics tools as every single short read carries little information by itself. In particular, repeat and paralogous sequences, as well as low-expressed regions and minor isoforms are difficult to assemble. Notice that transcripts that are moderately expressed only in a subpopulation of cells manifest an overall low expression level, as might be the case for long noncoding RNAs (lncRNAs) [4].

Unlike *de novo* transcript assembly approaches, which assemble reads solely based on the overlap of their sequences, genome-based methods employ a high-quality reference genome to better resolve ambiguities imposed by highly similar regions of the genome and to recover lower expressed transcripts. Genome-based methods first align reads to the genome to determine where each of the reads originated and then assemble the alignments into transcript models. This in turn introduces a critical dependence on the accuracy of the read alignment, which is affected by sequencing errors, polymorphisms, splicing, and ambiguous reads that belong to repeats. Reads spanning splice junctions between exons are particularly informative since they provide an explicit signal for the detection of splice donor and acceptor sites. At the same time, the spliced alignment of such reads is computationally challenging and error prone.

In case of an unbalanced split the prefix or suffix of a read that spans one of the two consecutive exons may be short and thus aligns equally well to a large number of genomic positions. Guessing the true origin can be further hampered by polymorphisms near the splice site. Besides *incorrect* spliced alignments this can also lead to *missed* splice junctions, i.e. exon-exon junctions that are not supported (covered) by any spliced alignment. Missed junctions can also result from read coverage fluctuations (biases) or a generally low transcript abundance. While some of the existing methods do take into account incorrect alignments by applying ad-hoc filters (Scripture [9], CLIIQ [10]) or by not requiring the isoform selection model to explain all input alignments (MITIE [11]), none of the existing approaches is able to deal with missed junctions. In this work we present a novel framework CIDANE (*C*omprehensive *I*soform *D*iscovery and *A*bu*N*dance *E*stimation), which, for the first time, allows to recover isoforms with uncovered splice junctions that are invisble to all existing approaches.

On a high level, existing methods for genome-based transcript assembly adhere to the following scheme: First, a set of candidate isoforms is defined as paths in a graph representing the base or exon connectivity as indicated by the aligned reads. Then, a *small* subset of isoforms is selected that explains the read alignments *well.* Since only a small number of transcripts is typically expressed in a given cell type (compared to the number of candidates), the restriction to few isoforms prevents fitting noise in the data.

Current methods mostly differ in the trade-offs they apply between the complexity of the model and the tractability of the resulting optimization problem, which largely determines the quality of the prediction: (i) Since the number of potential isoforms grows exponentially with the number of exons of the locus, all existing methods restrict either implicitly or explicitly the number of candidates they consider. Methods that do not enumerate isoforms explicitly either employ a simplified model with transcript-independent coefficients (e.g. MITIE [11], Traph [13]), or separate the intrinsically interdependent minimality and accuracy objectives (Cufflinks [2]). (ii) A second crucial algorithmic design decision is how to *balance* the two concurrent objectives. In an extreme case, the two objectives are treated independently (e.g., Cufflinks [2], CLASS [14], CLIIQ [10], Traph [13], IsoInfer [15]). More recent state-of-the-art methods (e.g., MITIE [11], iReckon [16], SLIDE [17], IsoLasso [18]) have recognize the importance of optimizing both objectives simultaneously and balance minimality and accuracy heuristically. (iii) Among methods that simultaneously optimize for both objectives, the measure of minimality has an enormous impact on the tractability of the resulting problem. The most immediate measure, the number of predicted transcripts (*L*^0^ norm), leads to non-convex objectives and a computationally intractable optimization problem. Methods like MITIE, Montebello [19], and iReckon, which employs a novel non-convex minimality measure, therefore resort to a forward stepwise regression strategy, a Monte Carlo simulation, or numerical optimization combined with random restarts, that generally do not find the best solution in this model. Methods like SLIDE and IsoLasso thus replace the *L*^0^ norm by the convex *L*^1^ norm, i.e. the sum of transcript abundances. (iv) Concerning the measure of accuracy, methods either apply a least-squares loss function (e.g. IsoLasso, SLIDE, TRAPH) or compute more generally a maximum likelihood assignment of reads to candidate isoforms. The latter typically requires a preselection of transcripts (Cufflinks) or lead to the intractability of the resulting problem (iReckon, Montebello).

Here we present CIDANE, a comprehensive tool for genome-based assembly and quantification of transcripts from RNA-seq experiments. The central idea of CIDANE is to trade the ability to determine the provably best transcript prediction in the underlying model for a slight approximation of the loss function. Intuitively, the accuracy and minimality measure (see (iii)-(iv)) fit noisy observations (read alignments) and thus the impact of their (adjustable) approximation on the overall prediction performance is expected to be rather limited. CIDANE therefore minimizes a least-squares loss function based on full-length transcripts and replaces the *L*^0^ minimality measure by the convex *L*^1^-norm. The *L*^1^ norm in fact *selects* a subset of transcripts with non-zero expression levels that is predicted to be expressed in a given cell type. A formulation based on full-length isoforms enables us to develop a comprehensive linear model (similar to SLIDE [17]) which, among others, takes into account the dependence of the distribution of read pairs along a given transcript on the estimated fragment length distribution. In contrast to previous methods, we employ a state-of-the-art machine learning algorithm to compute the optimal (according to a strict mathematical measure) balance between accuracy and minimality at essentially no additional computational cost. In a second phase, CIDANE linearizes the least-squares loss function with bounded error, which allows us to formulate our model based on all possible candidate transcripts without having to enumerate them explicitly. Following the principle of *delayed column generation* [20], we only add isoforms to our model “on demand”, i.e. if they help to strictly improve the overall prediction.

In contrast to Cufflinks, CIDANE implements a design that separates the assembly of full-length transcripts from the identification of its elementary components, i.e. exons or retained introns. This separation facilitates the incorporation of novel methods for splice site detection as well as additional sources of information to yield more accurate transcript assemblies. Not only a growing annotation of known splice sites, exon junctions, transcription start and end sites or even full-length isoforms can guide the assembly for most model organisms, but also additional gene boundary data can aid the interpretation of RNA-seq data. Our experiments demonstrate CIDANE’s superior performance in all these different scenarios of optionally available levels of annotation as well as in the interpretation of additionally available gene boundary data. Inferring transcript boundaries from RNA-seq read coverage drops is hampered by biases in the assay and is thus error-prone. We show that CIDANE’s integrated analysis of RNA-seq reads and reads obtained from the 5’ ends and polyadenylation sites of mRNA yields considerably more precise predictions of full-length transcripts than an interpretation of RNA-seq data alone. The general workflow of CIDANE is illustrated in Figure 1.

**Figure 1:**
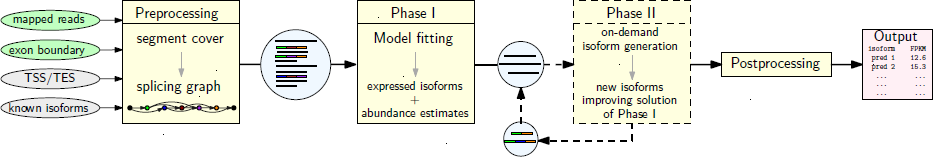
General workflow of CIDANE. Mandatory inputs (mapped RNA-Seq reads, exon boundaries) and optional inputs (transcription start (TSS) and end sites (TES), known transcripts) are used to summarize read alignments into segment covers, which count reads falling into non-ambiguously spliced segments of genes. From the corresponding splicing graph representation [12], an initial set of candidate isoforms is derived and a subset of expressed isoforms with estimated abundances is predicted by a regularized regression method during Phase I. This set forms the input to the optional Phase II, where improving isoforms are built on-demand by a delayed column generation approach. New candidates inferred in Phase II are then added to the initial candidate set to achieve a better fit of the model. After re-estimation of abundances and filtering (post-processing) a list of isoforms with abundance estimates is returned in gtf format.

## 2 Results and Discussion

We compared the performance of CIDANE in reconstructing transcripts from RNA-seq data to existing state-of-the-art methods. We evaluated the prediction quality on the transcript level based on both simulated and real data. While simulated data capture the characteristics of real data only to the extent that we understand the specifics of the experimental protocol, the performance analysis based on real RNA-seq data today still lacks a gold standard RNA-seq library along with annotated expressed transcripts. Therefore, the results of both types of experiments together provide a more meaningful picture of the true performance of a transcript assembly method.

Using simulated data, we investigated the impact of transcript abundance on the prediction quality and considered the scenario where a partial annotation of the (human) transcriptome is available to guide the reconstruction. We assessed both, the mere absence or presence of a (true) transcript in the prediction as well as the accuracy of the estimation of their abundances. Generating *perfect mapping files,* we make a first attempt to quantify the dependence of current genome-based transcript assembly tools on the accuracy of the read mapping. We demonstrate the superiority of CIDANE on real data through an integrated analysis of modENCODE RNA data, including RNA-seq, cap analysis of gene expression (CAGE), and Poly(A) Site sequencing (PAS-seq), obtained from heads of 20 day old adult *Drosophila melanogaster.* CAGE and PAS-seq data facilitate the mapping of transcription start and end sites, which are very difficult to infer from RNA-seq data alone.

In both cases, we compared the prediction to a reference transcriptome referred to as *ground truth* containing the *true transcripts.* Where not specified otherwise, we consider a true transcript as *recovered* by a predicted transcript if their intron sequences are identical. A true single-exon transcript is scored as recovered if it overlaps a predicted single-exon transcript. Every predicted transcript is matched to at most one true transcript and vice versa. If rec, true, and pred denote the number of recovered, true, and predicted transcripts, respectively, we applied recall 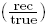, precision 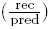, and F-score, the harmonic mean of recall and precision 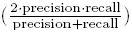, as a measure of prediction quality. To not penalize potential novel discoveries, the calculation of precision ignores predicted transcripts that do not overlap any of the reference transcripts.

### 2.1 Isoform reconstruction from simulated data

To obtain as realistic data as possible, we used the FluxSimulator [21] to generate RNA-seq datasets based on ∼78.000 UCSC-known (Feb. (GRCh37/hg19)) human transcripts [22]. After assigning randomized expression levels to all annotated transcripts following a distribution observed in real data, the FluxSimulator simulates the individual steps of an RNA-seq experiment, including reverse transcription, fragmentation, size selection, and sequencing. The version number of each tool along with parameters used in our experiments are specified in Additional file 2, Section 1.

#### 2.1.1 Ab initio transcript assembly

Mimicking the characteristics of real RNA-seq data, we generated four datasets comprising 40 and 80 million read pairs of length 75bp and 100bp, respectively. The fragment lengths observed after gel electrophoresis are modeled by a normal distribution *N*(250, 25) for the 75bp reads and *N*(300, 30) for the 100bp reads.^1^ We mapped each set of paired-end reads to the set of known transcripts using TopHat2 [23]. We defined the ground truth as the set of all annotated (UCSC) transcripts for which at least one paired-end read has been produced.

We compared the performance of CIDANE to the transcriptome reconstruction quality of Cufflinks [2], CLASS [14], IsoLasso [18], SLIDE [17], and Mitie [11]. We did not include iReckon in this first benchmark as it requires all known transcription start and end sites, which, as shown by the experiments in Sections 2.1.2 and 2.2, provides a valuable guidance in transcript reconstruction. While IsoLasso, SLIDE and CIDANE employ known exon-intron boundaries, Cufflinks and CLASS do not allow for the incorporation of pre-computed or annotated splice sites. Cufflinks does accept annotated full-length transcripts [24], a scenario which we will investigate in Section 2.1.2. In this experiment, we disable CIDANE’s ability to re-combine acceptor and donor sites to form novel exons. Since exon boundary information could be used to infer the originating strand, in the following we apply strand unspecific evaluation criteria. To eliminate a potential source of inaccuracy prior to the reconstruction algorithm, we provided IsoLasso and SLIDE with the fragment length distribution parameters as estimated by Cufflinks.

Figure 2(a) and 2(b) plot for each tool *X* a point with coordinates (precision of *X*, recall of *X*) and shows F-score isolines. For the dataset comprising 40 million 75bp read pairs (Figure 2(a)), CIDANE reconstructed transcripts with a recall value of 54.4%, a more than 18% increase over the recall achieved by Cufflinks (45.9%) and CLASS (43.7%), and a ∼30% improvement over IsoLasso (41.7%). At the same time, CIDANE predicts transcripts with the highest precision (71.6%), similar to Cufflinks (71.4%). IsoLasso (65.4%) seems to suffer from a heuristic determination of the regularization penalty. SLIDE appears at the lower left corner of the plot with recall 42.5% and 41.9%. Note that Cufflinks and CLASS model the transcript reconstruction problem as a *covering* problem minimizing the number of transcripts required to explain the input read alignments qualitatively. Neglecting quantitative information at this stage, it is not surprising that the two methods yield rather conservative predictions. Sections 2.1.2 and 2.2 show that the superior performance of CIDANE compared to Cufflinks cannot be attributed (only) to the additional exon boundary information. When provided with the exact same partial annotation of transcripts (Section 2.1.2) or when exon boundaries are inferred from the read data alone (Section 2.1.2), CIDANE still outperforms all existing methods.

**Figure 2:**
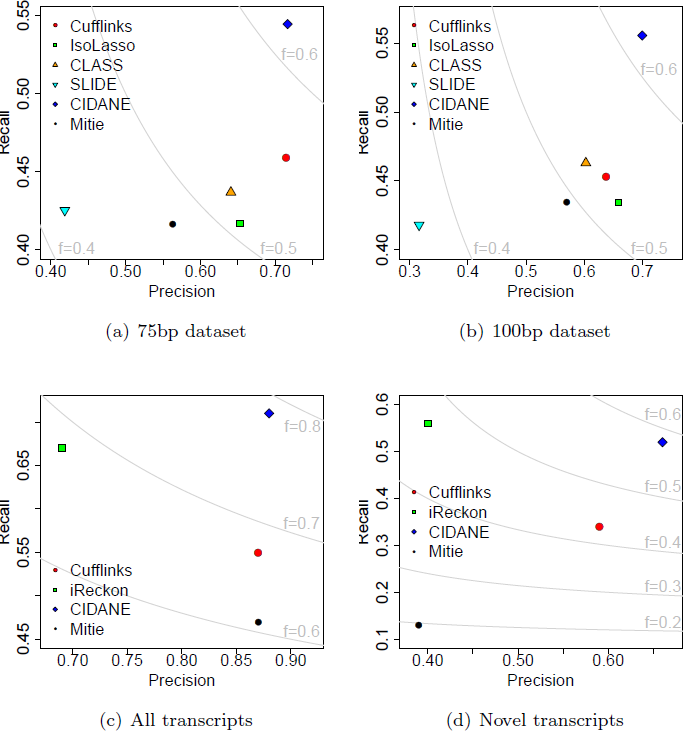
Each tool *X* ∊ {Cufflinks, IsoLasso, CLASS, SLIDE, CIDANE, Mitie} is represented by a point with coordinates (precision of *X*, recall of *X*). F-score isolines are shown in light-gray. Simulated datasets comprising 40 million 75bp (a) and 100bp read pairs (b), respectively. (c) and (d): Precision and recall achieved by each tool when provided a partial annotation.

The relative performance of the tools is similar when the same number of 100bp reads is generated (Figure 2(b)) or when the number of reads is doubled (Additional file 2, Figure 1). Cufflinks, however, seems to have difficulties assembling the 80 million 100bp read pairs. Recall and precision achieved by the tools for the four different experimental designs are listed in Additional file 2, Tables 1-4.

##### Dependence on transcript abundance

Further, we analyzed the influence of transcript abundance on the reconstruction capability of the different methods. We removed all transcripts that have most of their bases uncovered (< 0.1 FPKM) from the ground truth and split the remaining isoforms into three groups: *Low* comprises the 20% fraction of transcripts with lowest simulated expression, *High* the highest 5% fraction, and Med contains the remaining 75% of true transcripts. This subdivision corresponds to cutoffs in relative expression of ∼1.5 × 10^−6^ and ∼2.5 × 10^−4^ molecules, respectively. As expected, a higher abundance facilitates the reconstruction of isoforms (Figure 3). From the 75bp reads, however, CIDANE and SLIDE recover almost twice as many lowly expressed isoforms (recall ∼31% and ∼30%, respectively) as Cufflinks (recall ∼16.1%), whereas CLASS and IsoLasso recover only ∼6% and ∼3%, respectively. We observe similar results for the 100bp dataset. Not surprisingly, doubling the number of reads facilitates the recovery of low-expressed transcripts (Additional file 2, Figure 2). The ability of CIDANE to reconstruct, to some extent, even lowly expressed isoforms is likely due to its two core algorithmic improvements: First, CIDANE computes the entire regularization path in Phase I (see Section 4.1.2) to find the right balance between prediction accuracy and sparsity. An objective that is skewed towards sparsity typically yields predictions that miss low-expressed transcripts. Second, our approach considers a wider range of candidate transcripts than existing methods in Phase II (Figure 1). These include isoforms whose low abundance might cause splice junctions to be uncovered by reads rendering them invisible to other approaches. We investigate this effect in Section 2.1.4. Note that for the two 40 million read pairs datasets SLIDE achieves a similar recall on low-expressed isoforms only at the cost of a significantly lower precision and incurs a several orders of magnitude higher computational cost than CIDANE (see Section 2.1.5). When analyzing the two 80 million read pairs datasets, CIDANE reconstructs low-expressed transcripts with a ∼13% − 17% higher recall compared to SLIDE. All expression-level dependent recall values can be found in Additional file 2, Tables 1-4.

**Figure 3:**
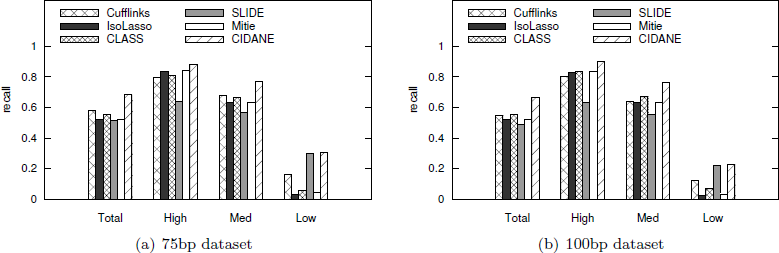
Recall achieved by the different methods in dependence of the expression level of true transcripts in the two 40 million read pairs datasets. Transcripts with simulated FPKM > 0.1 (*Total*) are grouped into sets *Low*, *High*, and *Med* which contain the lowest expressed 20% of the transcripts, the highest expressed 5%, and all remaining transcripts, respectively.

##### Dependence on mapping accuracy

In contrast to *de novo* assembly approaches, genome-based methods depend on the accuracy of the preceding mapping of reads to a reference genome. In this Section we make a first attempt to quantify this dependence. From reads simulated by the FluxSimulator [21], we generated a *perfect BAM file* that mapped each read to its true origin. In contrast, a BAM file output by TopHat2 or any other RNA-seq read aligner will generally contain incorrect mappings of reads caused by sequencing errors, polymorphisms, splicing, and read ambiguity due to repeats. We ran Cufflinks, IsoLasso, CLASS, Mitie, and CIDANE on the perfect BAM files of all four simulated data sets and compared recall and precision of the transcript assembly to the performance of the tools when provided with the BAM files generated by TopHat2 instead (Figure 4 and Additional file 2, Figure 3 and Tables 1-4). The difference in prediction accuracy will be a rather conservative estimate on the mapping dependency, since we expect a larger fraction of reads to be mapped incorrectly in real data than in idealized simulated data that neglect sequencing errors and certain types of biases. Nevertheless, when assembling the 40 million 75bp read pairs we observe a 1.1 – 2.4 and 2.0 – 11.7 percentage point improvement in recall and precision, respectively, when the true origin of all reads is known to the assembly tools (Figure 4(a)). Generally, assembly tools seem to benefit mostly in terms of precision rather than recall, independent of the experimental design. IsoLasso’s prediction precision seems to depend the most from perfectly aligned reads.

**Figure 4:**
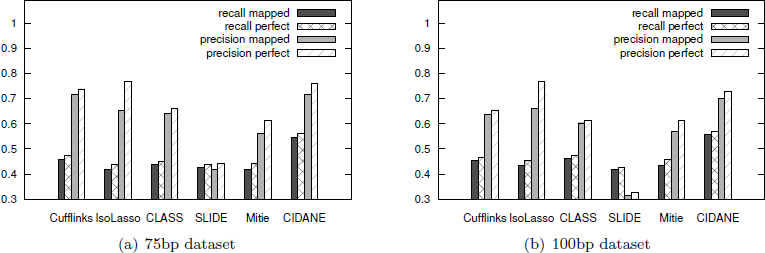
Dependence on mapping accuracy. Recall and precision of each tool is shown when provided a perfect BAM file (”perfect”) and when provided the mappings computed by TopHat2 (’’mapped”). Here we show the results on both 40 million read pairs datasets, similar results can be observed for the 80 million read pairs datasets (Additional file 2, Figure 3).

#### 2.1.2 Transcript assembly with partial annotation

We investigated the ability of Cufflinks, using the RABT approach presented in [24], iReckon, Mitie, and CIDANE to exploit an existing, but incomplete annotation of transcripts. No other assembly tool allowed to provide annotated transcripts. Such a partial annotation, available for the human transcriptome and many other studied organisms, can provide valuable guidance for the reconstruction of known isoforms, but algorithms must properly balance the preferential prediction of known transcripts and the detection of novel, unknown isoforms.

Our algorithmic scheme allows the incorporation of annotated transcription start and end sites during the isoform inference (see Sections 4.1.1 and 4.2.3). CIDANE accounts for a higher confidence in annotated versus novel transcripts by adjusted model parameters (see Sections 4.1.2 and 4.3).

From 1440 genes on chromosomes 1 and 2 for which between 2 and 8 isoforms have been annotated we randomly removed at least one and at most 50% of the known isoforms and provided each tool with the resulting ∼65% (Annot) of the originally ∼6300 known transcripts. The hidden ∼35% of annotated transcripts (New) constitutes the reference set (ground truth) in evaluating the ability of each method to infer novel isoforms in the presence of an incomplete annotation. As before, we used the FluxSimulator to generate 4 million read pairs (75bp), which were mapped to the ∼6300 transcripts by TopHat2 [23]. Experiments with 2 million and 8 million simulated read pairs led to similar performance results as shown below for 4 million read pairs.

Overall (Figure 2(c)), CIDANE predicted transcripts with highest recall and precision. CIDANE achieved a recall of ∼71.4%, which is ∼29% higher than the one of Cufflinks **(∼**55.3%), combined with a precision of ∼88%, which is ∼29% higher than the precision of iReckon (∼68%). Concerning the ability to correctly predict novel isoforms (Figure 2(d)), CIDANE recovered significantly more unknown transcripts than Cufflinks (recall ∼51.8% versus ∼34.1%). iReckon was able to reconstruct slightly more novel isoforms than CIDANE (∼55.8% versus ∼51.8%), but only at the cost of a lower recall with respect to annotated transcripts (Annot) (74% versus 82%) and a considerably lower precision (∼39.4% versus ∼65.7%). The precision in this case corresponds to the fraction of predicted novel isoforms matching an isoform in set New.

#### 2.1.3 Abundance estimation accuracy

In addition to evaluating the absence and presence of true transcripts in the prediction, we compared the accuracy of the abundance estimation of CIDANE to existing methods. We restrict this analysis to set Annot (see previous Section) to reduce the impact of isoform inference performance on the measure of abundance estimation quality. For every transcript in set Annot we compared the predicted FPKM (Fragments Per Kilobase of transcript per Million fragments sequenced) value to the true FPKM value calculated from the number of simulated paired-end reads. True transcripts that were not predicted by a method were considered as reconstructed with zero abundance. To reduce side effects on the abundance estimation due to very short transcripts, we limited the analysis to transcripts of length at least 500bp (∼98, 5%).

We observed similar Pearson correlation coefficients between true and predicted abundances for Cufflinks (0.96), iReckon (0.98), and CIDANE (0.97), see Fig. 5. To obtain a more detailed picture of the abundance estimation accuracy, we evaluated the relative error 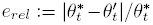 of reported abundance values 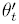 for transcript *t*, compared to the true abundances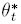.

**Figure 5:**
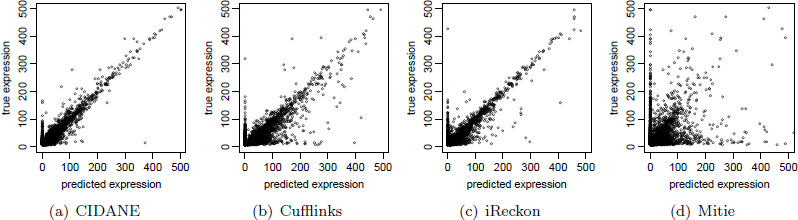
Correlation between simulated and predicted transcript abundance for FPKM values ≤ 500 on set Annot for (a) CIDANE, (b) Cufflinks, (c) iReckon, and (d) Mitie. Additional file 2, Figure 4 shows the full range of FPKM values, as well as log-scale plot after removing transcripts with predicted FPKM value 0.

Figure 6(a) plots the fraction of predicted transcripts with a relative abundance error below a certain threshold. Similarly, Figure 6(b) presents the recall values for each tool when scoring a true transcript as recovered only if the relative abundance error is below a certain threshold. Both Figures support the superior isoform reconstruction performance of CIDANE, not only in terms of correctly recovered isoforms, but also when taking into account their abundances. Isoforms with small relative error contribute predominantly to the improved performance of **CIDANE**, in particular compared to Cufflinks.

**Figure 6:**
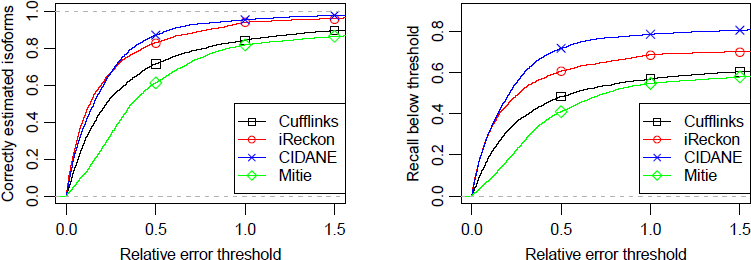
Relative error of predicted transcript abundances in FPKM. Left: Fraction of transcripts with non-zero predicted abundance implying a relative abundance error below a certain threshold. Right: Recall when scoring a transcript only if the relative abundance error is below a certain threshold.

#### 2.1.4 Delayed recovery of transcripts

In this benchmark, we demonstrate the capability of CIDANE to recover in Phase II (Figure 1) isoforms containing splice junctions that are not supported by any read. Note that a junction between neighboring exons can also be supported (”covered”) by a read pair which maps to the two exons, even if none of the reads spans the junction. From the (∼6300 transcripts expressed by the genes selected in the previous benchmark set, we simulated 2 million 75 bp read pairs. 118 transcripts had at least one splice junction uncovered and are therefore invisible to any method that derives candidate transcripts from a splicing graph representation of the read alignments (see Section 4.1.1). We note that this simulation neglects sequencing errors and any sequence specific or positional fragment biases. Furthermore, the mapping of reads to known transcripts is less error-prone than the spliced mapping to a reference genome and thus the number of such invisible isoforms is expected to be larger in practice. As before, CIDANE is given only the known exon boundaries and the mapped reads as input. For performance reasons, the delayed generation of transcripts was applied only to genes containing at most 50 exons, covering more than 99% of the genes. For larger genes CIDANE output the initial solution returned by our regularized linear regression approach (Phase I in Figure 1).

CIDANE successfully recovered (∼24.6% of the invisible transcripts expressed in our simulated cellular transcriptome. Cufflinks, Mitie, and IsoLasso (provided with exon boundaries) did not predict a single invisible isoform, while SLIDE recovered (∼5%. In rare cases, SLIDE in fact consideres candidates with uncovered junctions if otherwise only short candidates with at most 2 exons would exist (personal communication). We suspect that this strategy is one of the main causes for the very slow running time of SLIDE (see next Section).

When provided with a partial annotation (Annot) as in the previous benchmark, iReckon and Cufflinks recovered only one and two isoforms, respectively, whereas CIDANE recovered 17 ((∼40%) out of 42 invisible transcripts not contained in set Annot. Mitie again did not predict any invisible transcript. For each of the three invisible isoforms recovered by iReckon and Cufflinks, the provided annotation (Annot) reveals the uncovered splice junction within an alternative isoform. Neither of the methods was able to reconstruct any uncovered novel splice junction.

#### 2.1.5 Running times

All benchmarks were performed on a machine equipped with 2 Intel Xeon CPU X5550 @2.67GHz Quad Core and 72 GB memory, and all tools with multithreading support (Cufflinks, iReckon, SLIDE, Mitie) were allowed to use up to 16 threads. The current implementation of CIDANE uses only one thread. Only the pricing ILP solver uses up to 16 threads.

For the 75 bp dataset comprising 40 million read pairs assembled *ab initio* (Section 2.1.1), the running times of CIDANE in basic mode omitting Phase II (∼25 min), Cufflinks (∼21 min), IsoLasso (∼22 min), Mitie (∼40 min), and CLASS (∼73 min) were all within minutes to less than two hours, while SLIDE required ∼55.6 hours to complete. When assembling 80 million read pairs, the running time of CIDANE remains virtually unchanged, while it increases to ∼42 min for IsoLasso, ∼132 for CLASS, ∼59 min for Cufflinks, ∼125 min for Mitie, and to ∼62.3 hours for SLIDE. CIDANE’s optional search for invisible transcripts in Phase II requires an additional ∼31 min or ∼28 min of computation for the 40 and 80 million read pairs datasets, respectively. The number of reads mostly affects the preprocessing phase, where CIDANE counts reads mapping to unambiguous exon segments. The computational complexity of the optimization problems solved in Phase I and Phase II does not directly depend on the number of reads sequenced.

When providing an incomplete annotation (Section 2.1.2), we observed running times of ∼2 minutes for CIDANE, ∼3 minutes for Cufflinks, ∼5.5 min for Mitie, and ∼175 min for iReckon. Note that the latter experiment involves a smaller number of genes than the datasets analyzed in the *ab initio* setting.

### 2.2 Integrating real RNA-seq, CAGE, and PAS-seq

RNA-seq data provide an explicit signal for the detection of introns that is more informative than mere read coverage. Spliced alignments span splice junctions between exons and can be leveraged to infer splice donor and splice acceptor sites and thus the boundary of internal exons. In contrast, the reconstruction of transcript boundaries, i.e. the transcription start site (TSS) at the 5’ end and the transcription end site (TES) at the 3′ end, relies on a read coverage drop that is blurred by biases in the RNA-seq assay and is thus error-prone.

The conceptual separation of (i) the discovery of exons and (ii) the assembly of exons into transcripts allows CIDANE to employ additional sources of information in both modules. Not only a comprehensive (yet incomplete) annotation available for most model organisms can guide tasks (i) and (ii) (see Section 2.1.2), but additional gene boundary data can aid the interpretation of RNA-seq data [25].

By integrating *Drosophila melanogaster* RNA-seq, cap analysis of gene expression (CAGE), and Poly(A) Site sequencing (PAS-seq) data, GRIT [25] assembled transcripts with a considerably higher recall and precision than Cufflinks. CAGE and PAS-seq produce reads from the 5′ ends and polyadenylation sites of mRNAs, respectively, and thus facilitate the mapping of TSS and TES. Since reconstructing transcripts from RNA-seq data alone is intrinsically underdetermined [26], mapped TSS/TES can reduce the search space significantly, particularly for complex loci, and are thus expected to yield more accurate transcriptome predictions. In fact, experiments on simulated data performed in [15] already suggested the importance of TSS/TES information in transcript assembly.

In this section, we demonstrate the superiority of our comprehensive transcript assembly approach on the integrated analysis of modENCODE RNA data, comprising stranded RNA-seq, CAGE, and PAS-seq obtained from 20 days old adult *D. melanogaster* heads [25]. We reconstruct transcripts *ab initio* without relying on any elements of the annotation of the *D. melanogaster* genome. Instead, we employ exon and transcript boundary information obtained through the boundary discovery procedure of GRIT [25]: Exons and introns are identified by read coverage and spliced alignments, respectively; gene regions then contain exons that are connected by introns. In addition to splice donor and splice acceptor sites, TSS and TES are identified from read coverage peaks in the CAGE and PAS-seq data. For details we refer the interested reader to the original description of the procedure in [25].

Candidate transcripts considered by CIDANE correspond to paths in the splicing graph (see Section 4.1.1). In this experiment, CIDANE constructs nodes and edges representing exons and introns, respectively, that are discovered from the data by the above mentioned method. Only paths from exons whose 5′ boundary coincide with an identified TSS (and end with a splice donor site) to exons whose 3′ boundary coincide with an identified TES (and begin with a splice acceptor site) are considered. Single exon transcripts are bounded by identified TSS and TES on the 5′ and 3′ ends, respectively.

We compared the performance of CIDANE, GRIT (latest version 1.1.2c), and Cufflinks when reconstructing the cellular transcriptomes from stranded RNA-seq, CAGE, and PAS-seq generated from dissected heads of 20 days old adult *D. melanogaster*, using four replicates, two male and two female (see [25] or Additional file 2, Table 5). In the experiments performed in [25] on the same data sets, GRIT drastically outperformed annotation tools Scripture [9] and Trinity+Rsem[27] in terms of recall and precision. Here we apply the same evaluation criteria as in [25] and thus refrain from benchmarking CIDANE against tools Scripture and Trinity+Rsem. Similar to [25] we assumed a FlyBase 5.45 [28] transcript to be expressed in our sample if it is either composed of a single exon or if otherwise every splice junction is supported by at least one read. Since transcripts contained in the resulting ground truth by definition had no uncovered splice junctions, we disabled the delayed transcript recovery mode (Phase II in Figure 1) of CIDANE. Applying the above criteria, between ∼8200 and ∼10, 000 transcripts were expressed in each of the four *D. melanogaster* head samples.

We considered an expressed transcript in the resulting ground truth as successfully recovered if the sequence of introns (intron chain) match perfectly (same criteria as for simulated data) and if optionally the transcript boundaries, i.e. TSS and TES, lie within 50, 200, or 100, 000 bp of each other. Predicted transcripts that do not match any transcript in the ground truth but which are annotated in FlyBase do neither count as false positives nor as true positives. A comparison on the inton chain level ignores TSS and TES accuracy entirely and provides a meaningful measure of prediction quality if gene boundary information is either not available or not taken into account (Cufflinks). While a 50 and 200 bp tolerance assess the precision of transcript boundaries with different stringency, a relaxation to 100, 000 bp captures the multiplicities of identical intron chains that differ only by alternative promoters and/or polyadenylation sites.

Figure 7 depicts recall, precision, and F-score achieved by Cufflinks, GRIT, and CIDANE when predicting transcripts from integrated RNA data of adult *D. melanogaster* heads. Points of the same color correspond to the four replicates. Their precise coordinates are listed in Additional file 2, Tables 6-9. As was done in [25], we filtered transcripts predicted by GRIT with expression score lower bounds less than 1 × 10^−6^ estimated FPKM at a marginal 99% significance level.

**Figure 7:**
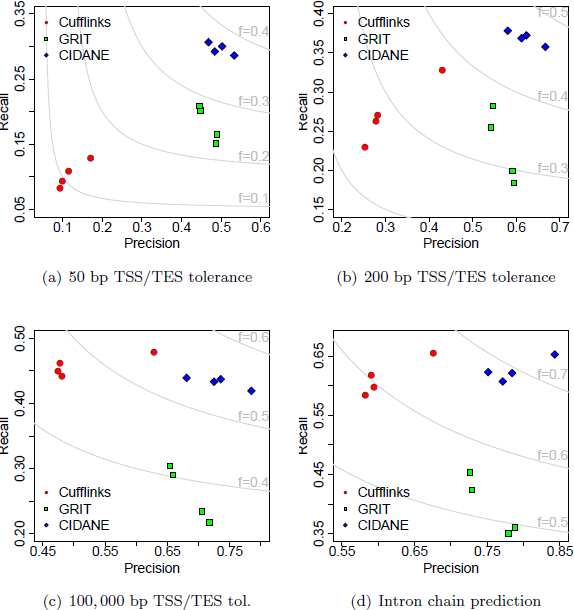
Recall and precision of transcript prediction by Cufflinks, GRIT, and CIDANE from integrated RNA data. Different thresholds in TSS/TES accuracy are applied in true positive definition.

Figures 7(a)-7(b) take into account the accuracy of transcript boundaries with different tolerances. If predicted and annotated TSS/TES are required to lie within 50bp of each other (Figure 7(a)), the lack of read data on the 5′ ends and polyadenylation sites of mRNAs results in a significantly poorer performance of Cufflinks compared to GRIT. Employing the same amount of data as GRIT, however, CIDANE achieves a recall of ∼29-31%, compared to ∼15-21% for GRIT, combined with a slightly higher precision. If we relax the TSS/TES tolerance to 200bp (Figure 7(b)), GRIT’s prediction profits from the additional CAGE and PAS-seq data mostly in terms of precision. Again, CIDANE manages to reconstruct more transcripts than GRIT with a slightly higher precision. Figures 7(c)-7(d) neglect the accuracy of transcript boundaries. In both cases, CIDANE combines the superior precision of GRIT with the superior recall of Cufflinks. Notice that in each analysis the transcriptome predictions of GRIT and CIDANE are based on the exact same mapping of exons, introns, TSS, and TES. The superiority of our approach results entirely from a more coherent assembly of exons into transcripts.

Concerning the efficiency, CIDANE ran less than two minutes per sample, while GRIT (allowing up to 16 threads) took ∼3h of computation. Note that GRIT’s runtime includes the discovery of exon and transcript boundaries.

## 3 Conclusion

We present CIDANE, which provides major improvements in cellular transcriptome reconstruction from RNA-seq over existing assembly tools. Through a carefully chosen trade-off between model complexity and tractability of the resulting optimization problem, and by applying state-of-the-art algorithmic techniques, CIDANE builds full-length transcript models from short sequencing reads with higher recall and precision than was possible before. CIDANE is engineered to not only assemble RNA-seq reads *ab initio*, but to also make use of the growing annotation of known splice sites, transcription start and end sites, or even full-length transcripts, available for most model organisms. Our experiments show that CIDANE’s core algorithmic engine yields more accurate transcriptome reconstructions than competing tools, in all these different scenarios and under various realistic experimental designs. Furthermore, CIDANE can employ additional gene boundary data to guide the assembly, thereby improving the precision of the reconstruction significantly.

To some extent, Phase II of CIDANE allows to recover splice junctions that are invisible to all existing approaches. Such junctions are not supported by any read alignment and can be observed predominantly among low-expressed transcripts. While CIDANE in basic mode (Phase II omitted) reconstructs a human cellular transcriptome from 80 million aligned read pairs in ∼25 min, the recovery of invisible junctions is a more complex task. For genes larger than 50 exons the iterative determination of invisible transcripts might become too expensive in practice and is disabled by default in our current implementation. Future work on the fixed-parameter tractability of the HEAVIEST ISOFORM problem might allow us to push the limits even further.

We expect that CIDANE will provide biologists with accurate transcript predictions from the very large, complex data sets that currently emerge from RNA-seq experiments. Such a high-resolution RNA-seq data interpretation is essential for any type of downstream analysis and will help to expand the catalogue of genes and their splice variants.

## 4 Methods

In this work, we assume mRNA fragments to be sequenced from both ends, yielding *paired-end reads.* Nonetheless, all results trivially apply to single-end reads. For each locus, identified as connected components of read mappings, CIDANE reconstructs isoforms from RNA-seq data in three phases (Figure 8). First (Section 4.1), a linear model is fitted (Figure 8c) to a compact representation of the observed read mappings (Figure 8a) using a set of fully supported candidate transcripts (Figure 8b). Here, our approach differs from existing methods mainly in (i) carefully designed regression coefficients that model (similar to SLIDE) the distribution of reads along a transcript, and in (ii) applying a state of the art machine learning algorithm to balance the accuracy of the prediction and the number of isoforms assigned a non-zero expression level. In a second phase (Section 4.2), CIDANE explores the space of transcripts that is neglected by existing methods due to computational challenges. To iteratively identify such a transcript that can help to improve the current prediction we have to solve a problem (Figure 8d) that we formalize as the HEAVIEST ISOFORM problem (HIS). If the “heaviest” isoform does not improve the current prediction CIDANE is guaranteed to have found the best possible set of isoforms without having explicitly enumerated all potential isoforms in the exponentially large space. Otherwise, the newly constructed isoform (Figure 8e) can be used to adjust our fitting.

**Figure 8:**
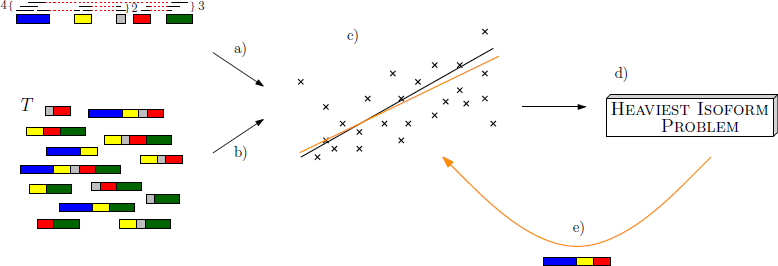
Schematic diagram illustrating CIDANE’s workflow. In the first phase, a linear model is fitted (black line in c) to a compact representation of the observed read mappings (a) using an initial set *T*_init_ of candidate transcripts (b). Second, transcripts not in *T*_init_ that can help to improve the prediction are iteratively identified as optimal solutions to the HEAVIEST ISOFORM problem (d). The newly constructed isoform (e) is used to adjust the fitting (orange line in c).

Although we show that HIS is NP-complete, we propose an integer linear programming (ILP) formulation that exploits certain properties of RNA-seq data and (optionally) known splicing characteristics that allow for the efficient solution of the ILP. For example, only a small number of combinations of exons enclosed by two mapped read mates is typically consistent with an estimated fragment length distribution, yielding a small number of variables in our formulation. Furthermore, we (optionally) disregard transcripts whose alternative promoter and polyadenylation sites coincide with acceptor and donor sites of internal exons, since signals read by the transcription and splicing mechanism to identify start (end) sites and acceptor (donor) sites differ significantly. Note that this restriction is conceptually equivalent to considering only *maximal* paths in the splicing graph as candidates, as is done by current methods. CIDANE, however, tries to restore maximal paths that are broken due to uncovered splice junctions. At the same time, the flexibility of an ILP formulation allows CIDANE to incorporate additional data or knowledge concerning, for instance, exon-intron boundaries, assembled exons, intron retentions, and transcription start and end sites.

The prediction is fine-tuned (Section 4.3) by re-fitting the linear model using the initial set of candidate transcripts augmented by all improving transcripts identified in the second phase of CIDANE. Finally, the expression levels of the reconstructed transcripts are re-estimated and converted into FPKM (Fragments Per Kilobase of transcript per Million fragments sequenced) in a post-processing phase (Section 4.3).

### 4.1 Phase I: Regularized linear regression

Similar to count based methods like SLIDE, IsoLasso, we summarize the observed read mappings into *segment covers* (Figure 8a). Instead of trying to explain each read mapping with it precise genomic coordinates we count the number of reads that fall into non-ambiguously connected segments of the genome. *Segments* in *𝒮* represent minimal exon fragments that are covered by reads and bounded by splice sites or transcription start or end sites (see Additional file 2, Figure 5), derived from spliced alignments, extracted from a set of gene annotations, or supported by additional data. For sequences of segments 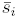 and 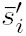, a *segment cover* 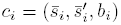 then counts the number *b*_*i*_ of read pairs *r* = (*r*_1_, *r*_2_) where *r*_1_ and *r*_2_ map with a *signature* consistent with 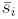 and 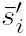, respectively, i.e. the mapping of *r*_1_ (*r*_2_) spans precisely the set of segment boundaries that are implied by 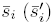 (see Additional file 2, Figure 6). *Faux segment covers* 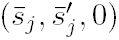 indicate that the corresponding combination of segments was *not* observed in the read data and can help to identify false positive predictions. We denote the set of segment covers, including faux covers (see Additional file 1, Section 1), by *𝒞*.

#### 4.1.1 Candidate Isoforms

We derive the initial set of candidate isoforms *T* (Figure 8b) used to explain the observations (segment covers) as paths in a *splicing graph* [12]. Nodes in a splicing graph correspond to segments *𝒮* and edges connect exon fragments whose consecutivity is indicated by (spliced) alignments. Under the assumption that every splice junction of every expressed isoform is covered by at least one mapped read, every expressed (true) transcript is among the paths in the splicing graph. For a formal specification of a splicing graph as employed in CIDANE see Additional file 1, Section 2.

We further define sets *𝒯𝒮𝒮* and *𝒫𝒜𝒮*, which contain potential transcription start and end sites, respectively. These sets can be compiled from annotated transcription start sites and polyadenylation sites, additional read data from the 5′ ends and polyadenylation sites of mRNAs (see Section 2.2), or purely from read mapping data. The latter is based on an exclusion principle: We do not allow for transcripts whose alternative promoter or polyadenylation sites coincide with acceptor and donor sites of internal exons and thus exclude all segments with spliced alignments supporting their 5′ or 3′ end from *𝒯𝒮𝒮* and *𝒫𝒜𝒮*, respectively. This exclusion strategy is equivalent to considering only *maximal* paths in the graph, as is done by current methods, and can easily be relaxed in CIDANE by setting *𝒯𝒮𝒮*: = *𝒮* and *𝒫𝒜𝒮*: = *𝒮*.

The set of candidate isoforms among which we select our initial prediction is then obtained by enumerating all paths in the splicing graph that start at a segment in *𝒯𝒮𝒮* and end at a segment in *𝒯𝒠𝒮*.

#### 4.1.2 Model fitting

We apply a linear model (Figure 8c) to estimate the number of reads originating from segments of the genome. Assuming that every position of an expressed transcript is equally likely chosen as starting position of a sequenced RNA fragment, we model the expected number of fragments mapping to segment cover *c =* (*s*, *s*′, *b*) as ∑_*t*∊*T*_ *ℓ*_*t,c*_*θ*_*t*_, where 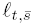 is the expected number of starting positions of fragments obtained from transcript *t* that show a mapping signature consistent with *c*. The expression level *θ*_*t*_ of transcript *t* counts the expected number of mapped fragments per transcript base (FPB), which is converted to FPKM at a later stage (Section 4.3). *ℓ*_*t,c*_ depends on the length of segments in 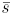 and 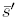, the length of segments in *t* enclosed by *s* and *s′*, the read length, and the cDNA fragment length distribution. Equations defining *ℓ*_*t,c*_ as used in our model are given in Additional file 1, Section 3. In contrast, methods like TRAPH, MITIE, and IsoInfer/IsoLasso define coefficients *ℓ*_*c*_ that neglect the dependence on transcripts *t*. Note that the distribution of reads along a transcript is generally not uniform, but typically unknown. The same applies to all the experimental data used in this study. Any prior knowledge concerning the likelihood of starting positions can be incorporated into our model through adjusted *ℓ*_*t,c*_ coefficients.

We employ the sum of squared errors (i.e. differences between estimated and observed number of reads) as a measure of accuracy of our prediction, weighted by an estimator for the variance of observations b [15]. Fitting our model using all candidate transcripts would allow to fit noise in the data by predicting a large number of isoforms with low but non-zero expression levels. Since in a given cell type really only a small subset of candidate transcripts is expressed our approach seeks a sparse set of expressed isoforms by augmenting, similar to SLIDE and IsoLasso, the objective by the *L*^1^-norm of the isoform abundances. Our (initial) prediction *θ* ≥ 0 comprises all transcripts with non-zero expression level in the optimal solution to:

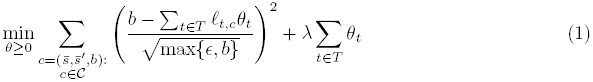

For faux covers, we replace *b* = 0 by ∈ (default: ∊ = 1). This so-called *Lasso* regression *selects* isoforms by setting the expression levels of all other transcripts to zero one at a time with increasing penalty terms λ.

The overall quality of the prediction crucially depends on the right choice of the regularization parameter λ. In contrast to previous methods, we balance the relative importance of the accuracy of the prediction and its simplicity (number of transcripts with non-zero expression level) based on the entire path of values for λ. As the coefficient path is piecewise linear, the entire regularization path can be computed at the cost of a single least-squares fit [29]. We apply a coordinate descent algorithm implemented in the `glmnet` Fortran code [30], that cyclically optimizes, for a given λ, each isoform abundance separately, holding all other abundances fixed. Update operations (inner products) directly profit from our sparse matrix of *ℓ*_*t,c*_ values (see Additional file 1, Section 3). Furthermore, considering a sequence of decreasing values for λ exploits estimates at previous λ’s as warm-start. After having computed the entire path of values for λ, our initial prediction is obtained from the optimal solution to (1) for the value of λ that yields the best *adjusted R*^2^ score. The adjusted *R*^2^ adjusts the goodness of fit (*R*^2^) for the number of isoforms used. If CIDANE is provided with a partial annotation of the transcriptome of an organism, the higher confidence in annotated transcripts is modeled by scaling the regularization penalties λ assigned to unknown transcripts by a factor of γ (default: γ = 2).

### 4.2 Phase II: Delayed generation of improving isoforms

The aim of CIDANE’s second phase is to recover isoforms with uncovered splice junctions (’’invisible transcripts”) that are not included in the candidate set of the regularized least squares regression due to their possibly very large number. We employ a delayed column generation technique [20] to identify new candidate isoforms that improve the optimal solution of the regularized least squares regression without exhaustive enumeration of all possible candidates. Particularly suited for large-scale linear programs, we formulate a piecewise-linear approximation (Additional file 1, Section 4) of the following quadratic program that is equivalent to the regularized least squares objective function (1):

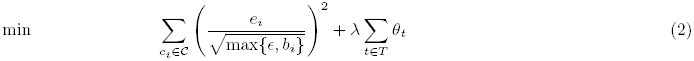

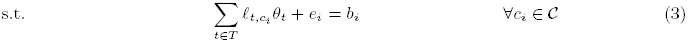

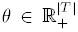 is the vector of transcript abundances, and *e* ∊ ℝ^|*c*|^ denotes the vector of errors, i.e. differences between estimated and observed read counts per segment cover. The generation of columns (i.e. variables *θ*_*t*_) is then accomplished by means of an ILP formulation presented below. In the following we let *m*: = |𝒞| be the number of segment covers falling into the considered locus and we let **A** be the corresponding coefficient matrix of constraints (3). Since the number of transcripts a gene can potentially encode grows exponentially with the number of its exons, constructing matrix **A** in full is impractical, even for comparatively small genes. Rather, we consider a restricted problem that contains only a small subset of all possible transcripts, represented by the *θ*-variables, and generate novel isoforms, i.e. columns of *A*, as needed to improve the overall prediction.

To identify an isoform that can help to improve the prediction in terms of objective (2), Dantzig’s simplex method [20] requires to determine a variable (transcript) *θ*_*t*__*j*_ with negative reduced cost 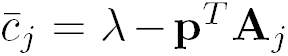 where **p** is the vector of simplex multipliers and **A**_*j*_ is the column of **A** representing transcript *t*_*j*_. Instead of computing the reduced cost associated with every possible transcript *t*_*j*_ we consider the problem of minimizing (λ – **p**^*T*^**A**_*j*_) over all *t*_*j*_, or equivalently, the problem of maximizing **p**^*T*^**A**_*j*_ over all transcripts *t*_*j*_. According to constraint (3), for every 1 ≤ *i* ≤ *m*, entry *i* of column **A**_*j*_ has value *ℓ*_*t*_*j*__,_*ci*_. The task is therefore to find a transcript *t*_*j*_ such that

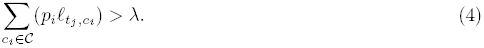

If no such transcript exists all reduced costs are nonnegative and the current solution is optimal. Next we model this optimization problem as a variant of the heaviest induced subgraph problem [31] and propose an integer linear programming formulation. For ease of notation, here we only consider the case where reads span single exons. For the general case of reads spanning an arbitrary number of exons we refer the reader to Additional file 1, Section 5.

Consider graph *G* = (*V*, *E*) that contains one vertex for each exon of a locus. We assume that the exons are numbered from left to right from 1 to n and identify each vertex by the corresponding exon number. We identify each segment cover 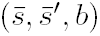 with single exon sequences 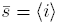, 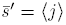 by (*i, j, b*) and include an edge e = (*i, j*) in *E*. For each edge *e* ∊ *E* we denote by 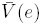 the set of vertices whose associated exons lie between the exons given by segments *i* and *j*, i.e. 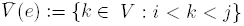. We assign to each edge *e* ∊ *E* a weight function 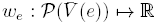. Then finding an improving transcript is equivalent to the following variant of the heaviest induced subgraph problem:

#### Definition 1

(HEAVIEST ISOFORM problem). *Given graph G* = (*V*, *E*) *and edge weight functions w*_*e*_, *find T* ⊆ *V such that the induced subgraph has maximal total edge weight, where each induced, edge e contributes weight*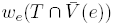

Edge weights *w*_*e*_ model the corresponding summands on the left-hand side of equation (4) and thus depend on the selection of exons between the mates of a cover (see Additional file 1, Section 3). In Additional file 1, Section 6, we show that the HEAVIEST ISOFORM problem is NP-complete. For single-end reads that span at most 2 exons the weight function is no longer dependent on 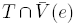 (e) and the HEAVIEST ISOFORM problem becomes polynomial-time solvable by a dynamic program.

This problem can be captured by the following integer linear program. For each vertex *i* in *G* a binary variable *x*_*i*_ indicates whether vertex *i* is contained in the solution. For every edge *e* ∊ *E* and every set 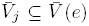 we have a binary variable *y*_*e,j*_ which is 1 if and only if vertices selected by the *x*-variables are consistent with 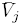 and induce *e*, enforced by the constraints below. In the objective function we let 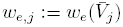:

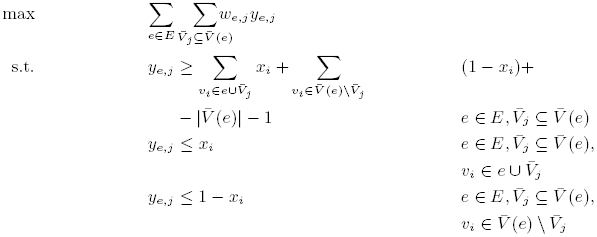

Depending on the quality of the data (determined by e.g. sequence specific or positional biases and read mapping accuracy) an isoform that is built by our ILP formulation might improve the prediction with respect to objective (1) by balancing, for instance, read coverage fluctuations. To prevent fitting noise in the data we require novel isoforms to explain segment covers c that are not supported by any transcript in the initial solution *T\** returned by the regularized least squares regression (1), i.e. ∀*t* ∊ *T*^*^:*ℓ*_*t,c*_ = 0. We refer to this set of initially unsupported segment covers as 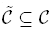. To reduce the impact of spurious read mappings we require a certain number *k*_*c*_ of read counts to be observed on the set of newly supported segment covers:

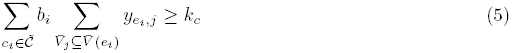

Intuitively, variables *y*_*e,l*_ associated with an edge *e* = (*i, j*) guess the selection of exons between exons *i* and *j*. Since for large *j* – *i* their exponential number would render our ILP approach infeasible, we neglect sets 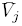 that would imply fragments of very unlikely length. More precisely, we apply lower and upper bounds 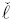 and 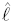 in the computation of ℓ_*t,c*_ (see equation (1) in Additional file 1) that limit the lower and upper 5%-quantile, respectively, of the estimated fragment length distribution. In Additional file 1, Section 7, we translate this fragment length restriction into lower and upper limits on the total length of exons in 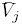, which allow us to enumerate feasible exon combinations in 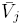 by an efficient splicing graph based backtracking scheme.

The construction of improving transcripts can be further guided by additional information such as exon-intron boundaries, transcription start and end sites, or exon connectivity. In the following we introduce constraints that we optionally add to our ILP formulation, depending on the available type of data, to ensure that the *x* variables encode a transcript that exhibits the desired structure.

#### 4.2.1 Exon compatibility

Splice acceptor and splice donor sites can be derived from spliced alignments or extracted from a set of gene annotations. Here we consider the case of a set of known exons *ε*. The more general case where the pairing of alternative acceptor and donor sites is unknown can be reduced to this case by simply including all possible combinations of acceptor and donor sites of an exon in *ε*. Alternatively, the structure of a splicing graph along with the individual mapping of acceptor and donor sites can be enforced through exon connectivity constraints as shown in the next Section.

To ensure that the segments in *𝒮* selected by the *x* variables form only valid exons in *ε* we link the segments of each exon *E*_*j*_ ∊ ε by an indicator variable *z*_*j*_:

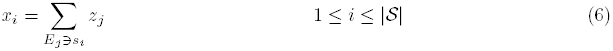

This constraint implies that (i) every selected segment i (i.e. *x*_*i*_ = 1) must be part of exactly one selected exon Ej (i.e. *z*_*j*_ = 1), (ii) all segments of a selected exon must be included, and (iii) no pair of overlapping, and hence incompatible, exons can be selected simultaneously.

#### 4.2.2 Exon connectivity

For some complexe genes it is computationally infeasible to enumerate all paths in the splicing graph to obtain the set of candidate isoforms. For such genes our delayed isoform generation approach allows the exploration of all candidate isoforms without explicitly enumerating them. Constraint (7) with *u*_*i,j*_: = 0 therefore captures the splicing graph structure in a way that the path induced by the selected set of segments agrees with the set of edges E in the splicing graph: A simultaneous selection of two segments s*i* and *s*_*j*_, *i < j*, without selecting any segment s_*k*_ with *i < k < j* is not feasible if the splicing graph does not contain edge (*v_i_,v_j_*). Notice that this scheme allows to assemble novel exons by selecting acceptor sites (incoming edge) and donor sites (outgoing edge) independently.

Alternatively, we can allow up to k (default: *k* = 2) new edges to be selected from a set of “valuable” edges E’ missing in the splicing graph. At most k binary variables *u*_*i,j*_, 1 < *i* < *j* < |*𝒮*|, can be set to 1 for (*v*_*i*_, *v*_*j*_) ∉ *E* to relax the corresponding constraint (7). We experimented with valuable sets of edges *E′* that allow the explanation of observed covers that cannot be explained using solely edges in *E*. In general, however, any novel intron can be simply modeled by a corresponding edge in *E′*.

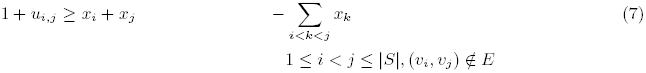

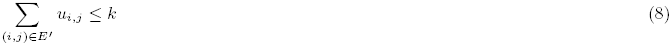

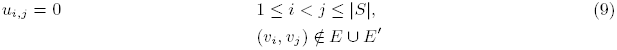

#### 4.2.3 Transcription start and end sites

We also have to ensure that improving transcripts built by our ILP start at segments in *𝒯𝒮𝒮* and end at segments in *𝒫𝒜𝒮*. Our model captures both the *exclusion* of potential transcription start and end sites from spliced alignments (see Section 4.1.1), and the *inclusion* of transcript boundaries, from e.g. a RNA-seq read coverage drop or from additional reads from the 5′ ends and polyadenylation sites of mRNAs (see Section 2.2).

Variables *ss*_*i*_ and *es*_*i*_ indicate the start and terminal segment of the generated isoform, respectively. We must select precisely one transcription start and end site (constraints (10)-(11)) from sets *𝒯𝒮𝒮* and *𝒯𝒠𝒮*, respectively (constraints (12)-(13)). Designated start and end sites must be part of the predicted transcript (constraint (14)-(15)). Finally, no segment upstream of the start segment (16) and no segment downstream of the end segment (17) can be part of the predicted isoform.

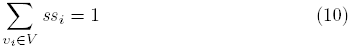

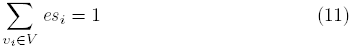

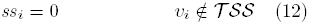

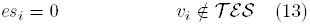

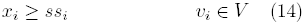

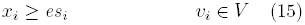

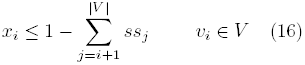

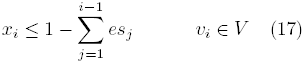

#### 4.2.4 Intron retentions

The explicit exon model described in Section 4.2.1 captures intron retentions by simply merging the flanking exons and the retained intron into one virtual exon that is added to set *ε*. Similarly, the more general exon connectivity formulation that is based on individual splice sites rather than assembled exons trivially includes the connectivity of intron retentions.

## 4.3 Phase III: Fine-tuning and post-processing

To adjust the regularization penalty λ to the increased set of candidate transcripts implicitly considered by the delayed isoform generation approach and to reduce the effect of the piecewise-linear approximation of the loss function, CIDANE resolves (1) with the candidate set T containing additionally all transcripts generated in the course of the delayed isoform generation phase. We increase the sensitivity of this fine-tuning step by reducing the regularization penalty prior to the delayed transcript generation by a multiplicative factor (default 0.9) and in turn express a higher confidence in fully supported isoforms by selectively increasing λ′ = α λ (default: α = 1.3) for delayed generated transcripts.

Let transcripts *T*^*^ = {*t*_1_,…, *t*_*m*_} with non-zero abundance 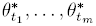 be returned by the regularized regression (1) solved in Phase I, optionally including the additional isoforms provided by our delayed isoform generation approach (Phase II). CIDANE determines the final prediction by post-processing *T*^*^ as follows. First, to avoid biases introduced by the regularization penalties λ we resolve (2)-(3) for λ: = 0 using set *T*^*^ instead of *T* to obtain expression levels 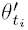. Second, we re-estimate the expression levels by computing a final assignment of mapped reads to isoforms that is guided by the relative abundances 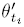:

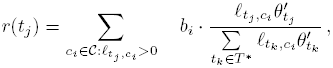

where *r*(*t*_*j*_) is the number of reads assigned to isoform *t*_*j*_. This assignment of reads to isoforms corrects overestimation or underestimation of the total number of reads within a gene due to nonuniform read mapping coverage. For all isoforms *t*_*j*_ ∊ *T*^*^ with *r*(*t*_*j*_) ≥ α (default: α = 10), we compute transcript expression levels in FPKM and finally return all isoforms whose predicted expression in FPKM is at least, *β*-percent (default: *β* = 10) of the expression of the most abundant transcript for the same gene. When run with a partial annotation of the transcriptome of an organism, we increase the expression threshold *β* to 20% for novel transcripts.

## Acknowledgements

We thank Nathan Boley for providing the data and support necessary for the integrated analysis of modENCODE RNA data. We thank Veronica Lee for her help in generating the perfect read mappings. We thank Sören Laue for helpful discussions on the computation of regularization paths. SC was supported in part by US National Institutes of Health grant R01-HG006677.

## Additional Files

**Additional file 1 — Algorithmic details**

**Additional file 2 — Additional figures and tables**

## Footnotes

1 The parameter files specifying the model of the RNA-seq experiment are available on our website http://ccb.jhu.edu/software/cidane/.

## References

[1] Pan, Q., Shai, O., Lee, L.J., Frey, B.J., Blencowe, B.J.: Deep surveying of alternative splicing complexity in the human transcriptome by high-throughput sequencing. Nat. Genet. 40(12), 1413–1415 (2008)

[2] Trapnell, C., Williams, B.A., Pertea, G., Mortazavi, A., Kwan, G., van Baren, M.J., Salzberg, S.L., Wold, B.J., Pachter, L.: Transcript assembly and quantification by RNA-Seq reveals unannotated transcripts and isoform switching during cell differentiation. Nature Biotechnology 28(5), 511–515 (2010)

[3] Wang, E.T., Sandberg, R., Luo, S., Khrebtukova, I., Zhang, L., Mayr, C., Kingsmore, S.F., Schroth, G.P., Burge, C.B.: Alternative isoform regulation in human tissue transcriptomes. Nature 456(7221), 470–476 (2008)

[4] Djebali, S., et al.: Landscape of transcription in human cells. Nature 489(7414), 101–108 (2012)

[5] Eswaran, J., Cyanam, D., Mudvari, P., Reddy, S.D., Pakala, S.B., Nair, S.S., Florea, L., Fuqua, S.A., Godbole, S., Kumar, R.: Transcriptomic landscape of breast cancers through mrna sequencing. Scientific Reports 2, 264 (2012)

[6] Seo, J.-S., Ju, Y.S., Lee, W.-C., Shin, J.-Y., Lee, J.K., Bleazard, T., Lee, J., Jung, Y.J., Kim, J.-O., Shin, J.-Y., Yu, S.-B., Kim, J., Lee, E.-R., Kang, C.-H., Park, I.-K., Rhee, H., Lee, S.-H., Kim, J.-I., Kang, J.-H., Kim, Y.T.: The transcriptional landscape and mutational profile of lung adenocarcinoma. Genome Res. 22(11), 2109–2119 (2012)

[7] Berger, M.F., et al.: Integrative analysis of the melanoma transcriptome. Genome Research 20(4), 413–427 (2010)

[8] Twine, N.A., Janitz, K., Wilkins, M.R., Janitz, M.: Whole transcriptome sequencing reveals gene expression and splicing differences in brain regions affected by alzheimer’s disease. PLoS ONE 6(1), 16266 (2011)

[9] Guttman, M., Garber, M., Levin, J.Z., Donaghey, J., Robinson, J., Adiconis, X., Fan, L., Koziol, M.J., Gnirke, A., Nusbaum, C., Rinn, J.L., Lander, E.S., Regev, A.: Ab initio reconstruction of cell type-specific transcriptomes in mouse reveals the conserved multi-exonic structure of lincRNAs. Nature Biotechnology 28(5), 503–10 (2010)

[10] Lin, Y.-Y., Dao, P., Hach, F., Bakhshi, M., Mo, F., Lapuk, A., Collins, C., Sahinalp, S.C.: Cliiq: Accurate comparative detection and quantification of expressed isoforms in a population. In: Raphael, B., Tang, J. (eds.) Algorithms in Bioinformatics. Lecture Notes in Computer Science, vol. 7534, pp. 178–189. Springer, Berlin, Heidelberg (2012)

[11] Behr, J., Kahles, A., Zhong, Y., Sreedharan, V.T., Drewe, P., Rätsch, G.: Mitie: Simultaneous rna-seq-based transcript identification and quantification in multiple samples. Bioinformatics 29(20), 2529–2538 (2013)

[12] Heber, S., Alekseyev, M., Sze, S.H., Tang, H., Pevzner, P.A.: Splicing graphs and EST assembly problem. Bioinformatics (Oxford, England) 18 Suppl 1, 181–188 (2002)

[13] Tomescu, A.I., Kuosmanen, A., Rizzi, R., Mäkinen, V.: A novel min-cost flow method for estimating transcript expression with RNA-Seq. BMC Bioinformatics 14 (suppl 5), 15 (2013)

[14] Song, L., Florea, L.: CLASS: constrained transcript assembly of RNA-Seq reads. BMC Bioinformatics 14 (suppl 5), 14 (2013)

[15] Feng, J., Li, W., Jiang, T.: Inference of isoforms from short sequence reads. J. Comput. Biol. 18(3), 305–321 (2011)

[16] Mezlini, A.M., Smith, E.J., Fiume, M., Buske, O., Savich, G.L., Shah, S., Aparicio, S., Chiang, D.Y., Goldenberg, A., Brudno, M.: iReckon: simultaneous isoform discovery and abundance estimation from RNA-Seq data. Genome research 23(3), 519–529 (2013)

[17] Li, J.J., Jiang, C.-R., Brown, J.B., Huang, H., Bickel, P.J.: Sparse linear modeling of next-generation mRNA sequencing (RNA-Seq) data for isoform discovery and abundance estimation. Proceedings of the National Academy of Sciences 108(50), 19867–19872 (2011)

[18] Li, W., Feng, J., Jiang, T.: IsoLasso: A LASSO regression approach to RNA-seq based transcriptome assembly. Journal of Computational Biology 18(11), 1693–1707 (2011)

[19] Hiller, D., Wong, W.H.: Simultaneous isoform discovery and quantification from RNA-seq. Statistics in Biosciences 5(1), 100–118 (2013)

[20] Bertsimas, D., Tsitsiklis, J.N.: Introduction to Linear Optimization. Athena Scientific, Belmont (MA) (1997)

[21] Griebel, T., Zacher, B., Ribeca, P., Raineri, E., Lacroix, V., Guigó, R., Sammeth, M.: Modelling and simulating generic rna-seq experiments with the flux simulator. Nucleic Acids Research 40(20), 10073–10083 (2012)

[22] Hsu, F., Kent, W.J., Clawson, H., Kuhn, R.M., Diekhans, M., Haussler, D.: The UCSC known genes. Bioinformatics 22(9), 1036–1046 (2006)

[23] Kim, D., Pertea, G., Trapnell, C., Pimentel, H., Kelley, R., Salzberg, S.: Tophat2: accurate alignment of transcriptomes in the presence of insertions, deletions and gene fusions. Genome Biology 14(4), 36 (2013)

[24] Roberts, A., Pimentel, H., Trapnell, C., Pachter, L.: Identification of novel transcripts in annotated genomes using RNA-Seq. Bioinformatics (2011)

[25] Boley, N., Stoiber, M.H., Booth, B.W., Wan, K.H., Hoskins, R.A., Bickel, P.J., Celniker, S.E., Brown, J.B.: Genome-guided transcript assembly by integrative analysis of RNA sequence data. Nat Biotech 32(4), 341–346 (2014)

[26] Lacroix, V., Sammeth, M., Guigo, R., Bergeron, A.: Exact transcriptome reconstruction from short sequence reads. In: Crandall, K.A., Lagergren, J. (eds.) Algorithms in Bioinformatics. Lecture Notes in Computer Science, vol. 5251, pp. 50–63. Springer, Berlin, Heidelberg (2008)

[27] Grabherr, M.G., Haas, B.J., Yassour, M., Levin, J.Z., Thompson, D.A., Amit, I., Adiconis, X., Fan, L., Raychowdhury, R., Zeng, Q., Chen, Z., Mauceli, E., Hacohen, N., Gnirke, A., Rhind, N., di Palma, F., Birren, B.W., Nusbaum, C., Lindblad-Toh, K., Friedman, N., Regev, A.: Full-length transcriptome assembly from RNA-Seq data without a reference genome. Nat Biotechnol 29(7), 644–652 (2011)

[28] Marygold, S.J., Leyland, P.C., Seal, R.L., Goodman, J.L., Thurmond, J., Strelets, V.B., Wilson, R.J., the FlyBase consortium: Flybase: improvements to the bibliography. Nucleic Acids Research 41(D1), 751–757 (2013)

[29] Efron, B., Hastie, T., Johnstone, I., Tibshirani, R.: Least angle regression. The Annals of Statistics 32(2), 407–499 (2004)

[30] Friedman, J., Hastie, T., Tibshirani, R.: Regularization paths for generalized linear models via coordinate descent. Journal of Statistical Software 33(1), 1–22 (2010)

[31] Kortsarz, G., Peleg, D.: On choosing a dense subgraph. In: Foundations of Computer Science, 1993. Proceedings., 34th Annual Symposium On, pp. 692–701 (1993)

